# Long-Range Hopping Conductivity in Proteins

**DOI:** 10.1101/2022.10.27.514097

**Authors:** Siddharth Krishnan, Aleksei Aksimentiev, Stuart Lindsay, Dmitry Matyushov

## Abstract

Single molecule measurements show that many proteins, lacking any redox cofactors, nonetheless exhibit electrical conductance on the order of a nanosiemen, implying that electrons can transit an entire protein in less than a nanosecond when subject to a potential difference of less than 1V. In the conventional fast transport scenario where the free energy barrier is zero, the hopping rate is determined by the reorganization energy of approximately 0.8 eV, which sets the time scale of a single hopping event to at least 1μs. Furthermore, the Fermi energies of metal electrodes used in experiments are far-removed from the equilibrium redox states of the aromatic residues of the protein, which should additionally slow down the electron transfer. Here, we combine all-atom molecular dynamics (MD) simulations of non-redox active proteins (consensus tetratricopeptide repeats) with an electron transfer theory to demonstrate a molecular mechanism that can account for the unexpectedly fast electron transfer. According to our MD simulations, the reorganization energy produced by the energy shift on charging (the Stokes shift) is close to the conventional value of 0.8 eV. However, the nonergodic sampling of molecular configurations by the protein results in reorganization energies, extracted directly from the distribution of the electrostatic energy fluctuations, that are only ~ 0.2 eV, which is small enough to enable long-range hopping. Using the MD values of the reorganization energies we calculate a current decay with distance that is in agreement with experiment.

**Significance:** Electron transfer is fundamental to biology, facilitating a range of metabolic processes and efficient energy conversion. Conventionally, electron transfer through proteins is thought to occur via a chain of metal or organic co-factors connecting one side of the protein to another. Recent experiments, however, show that proteins lacking any co-factors can nonetheless transport electrons with high efficiency if properly connected to metal electrodes. This study provides a theoretical model of such cofactor-less transfer, showing that transient occupation of non-equilibrium states of the protein’s aromatic residues reduces the barrier to electron hopping, facilitating long range and rapid transport. Our results widen the pool of proteins potentially involved in biological electron transport and provide theoretical underpinning to design of protein molecular electronics.

## Introduction

Proteins participating in the energy chains of biology (photosynthesis and respiration (1) and other enzymatic reactions) have to change the oxidation state of their active sites. Since amino acids are mostly redox inactive, the prevailing dogma in the field is that changes of the oxidation state are achieved by utilizing redox active cofactors intercalated into the protein fold. The possibility that this assumption is not always true has been considered in studies of electron relays in proteins (2–6) where protein residues effectively act as semiconductor elements to facilitate exchange of electrons between active sites. This conductivity mechanism adds versatility to redox active enzymes because direct tunneling in biology is limited to ≃ 1.4 nm (7). If protein residues conduct, can they do so on the time scales of an enzymatic turnover?

It is often suggested that the barrier required for the electron to reach the tunneling configuration to hop between the localization sites is described by Marcus theory (8) with a “universal” protein reorganization energy (7) *λ* ≃ 0.8 eV and a reaction free energy Δ*G*. The latter should be close to zero for hops between equal residues, in which case the barrier is mostly determined by the magnitude of *λ*. Some dependence of the tunneling probability on the bridging medium was found (9), but the experimental distinctions between intervening media are not yet sufficient to allow for selection of specific tunneling pathways through particular amino acid residues. These two components, a fairly large and uniform reorganization energy, and lack of knowledge of a specific tunneling path, are obstacles to calculating the conductivity occurring via hops between aromatic residues (mostly tryptophans, tyrosines, and histidines (5, 10)). In particular, the rates of individual hops are not sufficiently high to allow for a hopping conductivity of the magnitude observed experimentally (11–14).

The conductance of single protein molecules can be measured directly, provided that chemical contacts are utilized to inject charge (12), and a summary of one such set of measurements (13) for a series of linear proteins (the consensus tetratricopeptide repeat proteins – CTPR) is given in Figure 1. These proteins consist of a two-helix motif that is readily concatenated via recombinant techniques to form oligomers of a controlled length. The measured single molecule resistance is ~ 1GΩ over a distance of 15 nm, fairly typical of many non-redox active proteins (12). This equates to a current of 0.1 nA at a bias of 0.1V, or about one electron passing from one contact to the other every ns. In contrast, an estimate (c.f. Equation 5) of the hopping time for a characteristic 0.6 nm distance with λ = 0.8 eV yields a time between hops of ~ 1 μs, three orders of magnitude too slow. Eshel et al. (15) have pointed out that similar problems arise in an analysis of OmcS bacterial wires – specifically that in order to account for the measured conductivity, a reorganization energy below 0.2 eV and a fast diffusion constant (~ 20 nm^2^/ns) are required.

**Figure 1:**
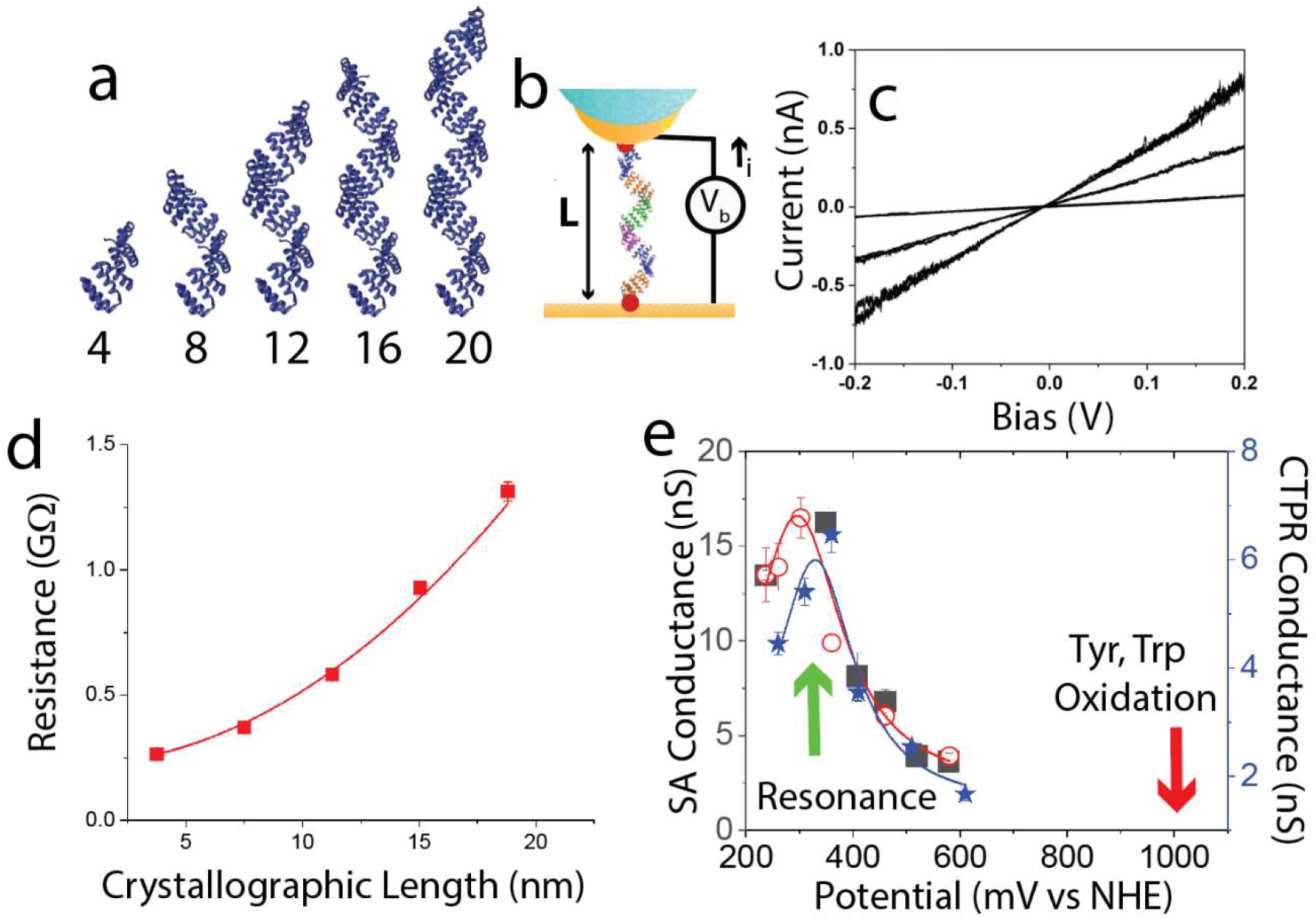
Experimental single-molecule conductance as a function of length of linear proteins: **a)** Five consensus teratricopeptide repeat (CTPR) proteins differing by the number of the repeat units as listed below the images. **b)** Schematic of a scanning tunneling microscope (STM) measurement of protein conductance. Cysteines (red dots) at the N- and C-termini form contacts between the protein and the metal electrodes. **c)** Electrical current versus electrode voltage difference. Examples of recordings from 3 different CTPR8 molecules are shown. Many such recordings yield a distribution of single-molecule resistances. **d)** The most probable resistance of a CTRP protein versus its length. The resistance does not increase linearly with L but rather as L^2^ (red line) with an intercept of 0.22 GΩ. (data from ref. 12) **e)** The molecular conductance depends on charge injection potential, as controlled by the Fermi energies of the metals used (black squares) or by changing surface polarization under potential control (open circles) – data (from ref. 15) are for streptavidin (SA) in both cases. Stars are data for CTPR8, the protein considered here (data from ref. 12). For both proteins the conductance has a maximum at about 300 mV vs. the normal hydrogen electrode, NHE, (green arrow), well removed from the oxidation potentials of tyrosine and tryptophan at ~1V vs. NHE (red arrow).

A second problem lies in the electrochemical potential dependence of the conductance (16). This is measured either by changing electrode materials (to change the Fermi energy) or by changing the surface polarization of a given metal under electrochemical potential control (16). The conductance measured by either method shows a sharp maximum at ~ 300 mV vs. the normal hydrogen electrode, NHE, (Figure 1e). This is quite different from the ~1V vs NHE required to oxidize tyrosine and tryptophan (17, 18). As we shall show, these challenges are largely resolved with the appropriate treatment of the reorganization energy.

Below, we describe the required modification to Marcus theory, then use all-atom molecular dynamics (MD) simulations to calculate the effective reorganization energy for charge transfers among the aromatic residues of CTPR monomers and dimers, and calculate the relative displacements of these residues, in order to calculate the hopping rates. A kinetic Monte Carlo simulation of the carrier diffusion, taken together with a model for charge injection, predicts both the magnitude of the currents and their decay with distance in agreement with experiment.

### Modifications to Marcus Theory

Here, we use the approach of Warshel (19) who amended the Marcus picture of crossing parabolas by specifying the energy gap *X* between the donor and acceptor energies of the electron as the reaction coordinate. The probability of reaching the tunneling configuration, *X* = 0, is given by Gaussian statistics, as a consequence of long-range electrostatics of the electron interacting with many particles in the medium (*i.e*., the central limit theorem). The combination of the Gaussian distribution of the reaction coordinate *X* with the fluctuation-dissipation theorem (FDT) (20, 21) leads to a specific connection between the separation of distribution maxima in the initial and final states (the Stokes-shift reorganization energy, *λ*^st^ (22)) and the Gaussian width, 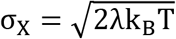, specified by the Marcus reorganization energy *λ*. The Marcus theory utilizes this connection to establish a single reorganization energy for electron transfer λ^St^ = *λ*. This simplification leads to the activation barrier for hopping electron transfer Δ*G^†^* in terms of the two parameters, the reorganization energy *λ* and the reaction free energy Δ*G*:

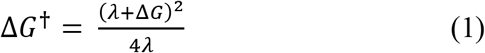

While the Gaussian distribution of the energy gap is a general consequence of statistics of many interacting particles, the application of the FDT to the statistics of the energy gap requires a Gibbsian ensemble based on the assumption that all “essential” configurations are accessible by the protein thermal bath on the reaction time scale. This is often not the case for a solvated protein, as the thermal bath, and many configurations are not accessible, either because they require times that are too long to be reached, or they are thermodynamically unstable (e.g., protein unfolding) or geometrically prohibited by closely packed folded protein. Overall, a folded protein is a frustrated (glassy (23)) medium with many dynamical and geometrical constraints imposed on the configurations coupled to the reaction coordinate. These frustrations lead to a non-Gibbsian statistics of thermal fluctuations and to violations of the FDT (22). When sampling is incomplete (nonergodic), the direct correspondence between *λ*^st^ and *λ* is broken. In particular, the parabolas connecting the free energy distributions to *X* can be broader (solid lines in Figure 2a) than predicted by Marcus theory (dashed lines in Figure 2a), leading to an effective (reaction) reorganization energy (22, 24)

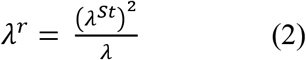

that replaces the Marcus *λ* in Eq. 1. Simulations of electron transfer between redox cofactors in electron-transfer proteins showed *λ*^st^ < *λ* and thus there is a lower barrier for hopping electron transfer than suggested in the Marcus framework (22, 24, 25). Low activation barriers achieved through λ^*r*^ < λ can potentially lead to sufficiently fast hops between aromatic residues, thus forming conductivity relays for protein electron transfer (3, 4). However, the required inequality λ^st^ < *λ* has been established only for protein cofactors (22), and it remains unclear if the inequality also applies to hopping conductivity through aromatic residues. According to our MD simulations (described below), a significant reduction of the activation barrier is indeed achieved for electron hops between tyrosine and tryptophan residues due to 〈*λ*^st^/*λ*〉 ≈ 0.4. This ratio also enters the nonergodic reaction free energy which is modified from the thermodynamic value Δ*G*_0_ to

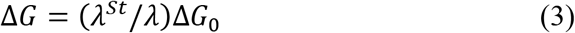

**Figure 2:**
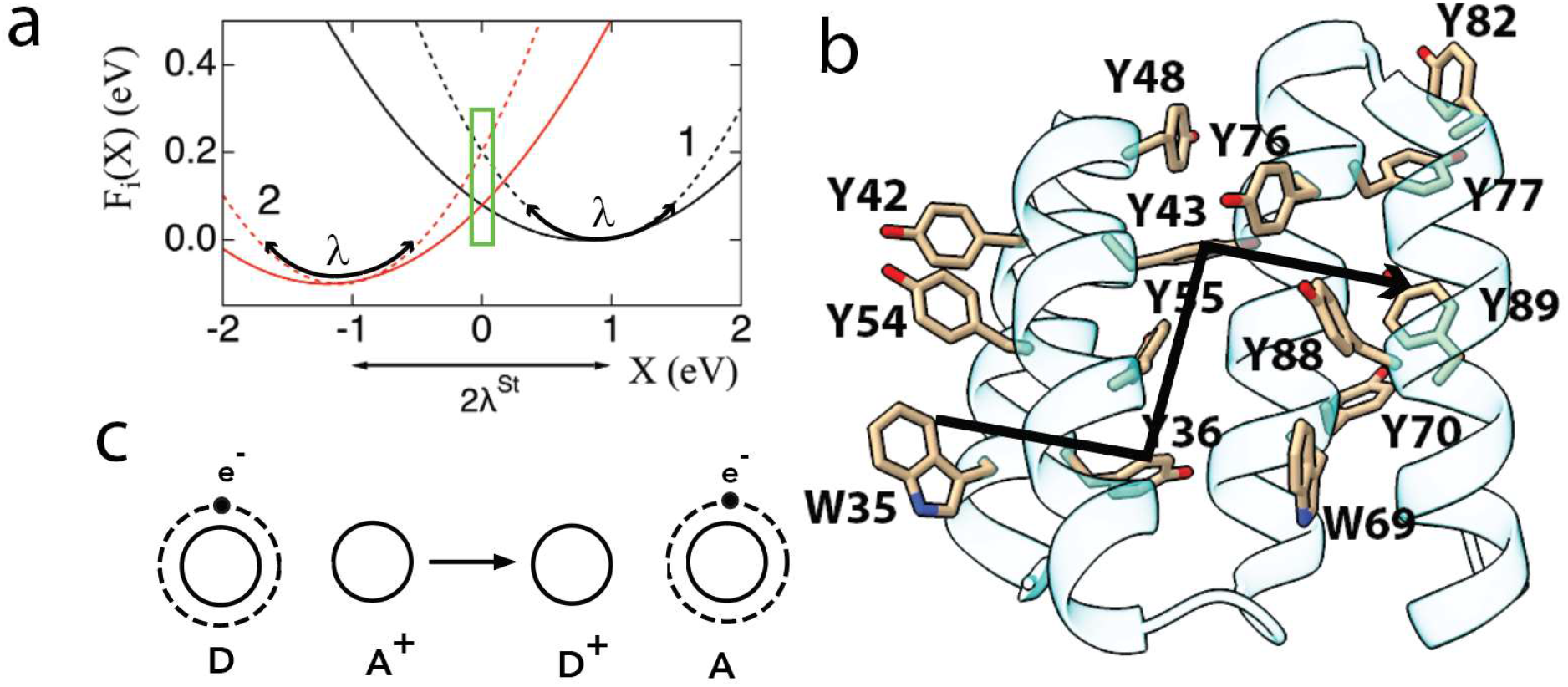
The hopping mechanism: **a)** Schematic free energy diagram of a donor (D)--acceptor (A) pair as a function of the electrostatic potential difference *X* between the states before (“1”) and after (“2”) a charge transfer. The potential difference between the free energy minima is twice the Stokes shift for optical transitions, λ^St^. Tunneling transitions occur where the parabolas intersect (green box). In Marcus theory, λ^St^ also determines the width of the parabolas (dashed lines) through the fluctuation-dissipation theorem. However, when electron transfer is fast compared to the thermal equilibration time, the width of the parabolas (solid lines) is no longer determined by λ^St^ and must be explicitly calculated from the distribution of energy fluctuations, leading to a reduced reorganization energy, λ^*r*^, with a corresponding reduction in the barrier for electron transfer (compare the crossing point of the solid parabolas vs. the crossing point of the dashed parabolas). **b)** Hopping occurs via aromatic residues. Two repeat units of CTPR protein are shown with tyrosines and tryptophans highlighted (numbering includes an N-terminal tail). The arrow shows one of the many paths through the two turns of protein, with W35 as the entry point and Y89 as the exit. **c)** Residues participating in the charge shift reaction change their oxidation state as illustrated (D = electron donor, A = electron acceptor).

Taking the ratio of the reorganization energies into account, the nonequilibrium redox potential of the tyrosines and tryptophans approaches the experimentally determine value of ~300 mV vs. NHE, Figure 1e.

The next important question is how individual random hops of charge carriers combine to generate the overall current and how the protein conductivity scales with the distance between the points of charge injection connected to external electrodes. The experimental data (13) summarized in Figure 1 (and data from other experiments (26)) show a very slow decrease of conductivity with the size of the protein. Previous studies suggested that this weak scaling reflects coherent tunneling of electrons through the entire protein (27, 28). Existing theories and calculations do not support this hypothesis (29) prompting the need for an alternative conductance mechanism.

Here, we propose that protein conductivity arises from random hopping of charge carriers over the network of tyrosine and tryptophan (Figure 2b) sites of the protein upon injection of the charge carriers at the electrode contact sites. The conductivity of redox-inactive proteins is mostly determined by the chemistry of the contact (12) and the externally imposed bias is dropped largely at the contacts. The random walk does not involve a significant external driving force, so the dependence of conductivity on the protein size is determined by the hopping sites involved and the hopping rates. We develop a quantitative theory using all-atom simulations to calculate the electrostatic energy fluctuations of donor and acceptor sites in the initial and final states, thus deriving the effective reorganization energies and the free energies for the various hops between multiple pairs of aromatic residues located within the Dutton radius (7) (1.4 nm). The effective reorganization energies are used to calculate the hopping rates, from which a diffusion constant is derived, via a set of Monte Carlo simulations, leading to calculation of the experimentally-observed quantities.

### Extracting Local Hopping Rates from Molecular Dynamics Simulations

A CTPR8 protein contains a total of 60 aromatic residues, out of which 52 are tyrosine and eight are tryptophan. The protein consists of eight two-helix repeat units with each unit being 34 amino acids in length and containing six tyrosines and one tryptophan (two such units are shown in Figure 2b). We hypothesized that, during the charge transfer, these residues are oxidized to their cationic free radical states (Figure 2c). We probed this possibility by conducting all-atom MD simulations of the protein having the aromatic residues in their neutral and oxidized states and using the fluctuations of the residues’ electrostatic energy to compute the rate of inter-residue electron transfer.

Figures 3a and b illustrate two all-atom systems used to probe a transfer of electron from residue Y77 to Y43. In the initial state of the hopping process (Figure 3a), Y43 is in its oxidized state whereas Y77 is charge neutral. At the end of the hopping process, Y77 is oxidized and Y43 is neutral. To evaluate the rate of electron transfer we computed equilibrium fluctuations of the electrostatic energy gap *X*(*t*) = ∑_*i*_ Δ*q*_*i*_φ_*i*_, where the sum runs over all atoms of the two residues accepting or donating an electron, Δ*q*_*i*_ = *q*_*i*_^final^ − *q*_*i*_^initial^ is the change of the partial charge of atom *i* upon transfer of the electron, and Φ_*i*_ is the local electrostatic potential at atom *i*. Knowing the partial charges on the oxidized forms of a tyrosine (30) and a tryptophan (31) residue (Tables S4 and S5), we determined the energy gap fluctuations by performing a 0.75 microsecond equilibration of each system and evaluating the instantaneous distribution of the electrostatic potential every 10 ps using the PMEpot plug-in of VMD (32, 33). Figure 3c plots the resulting *X*(*t*) data for the initial and the final states of the Y77-Y43 transfer, whereas Figure 3d shows their statistical distributions. The corresponding free-energy distributions of the Y77-Y43 pair, obtained from the electrostatic energy gap statistics as *G*_*i*_ = −*k*_B_*T*ln[*P*_*i*_(*X*)], are shown in Figure 3e.

**Figure 3:**
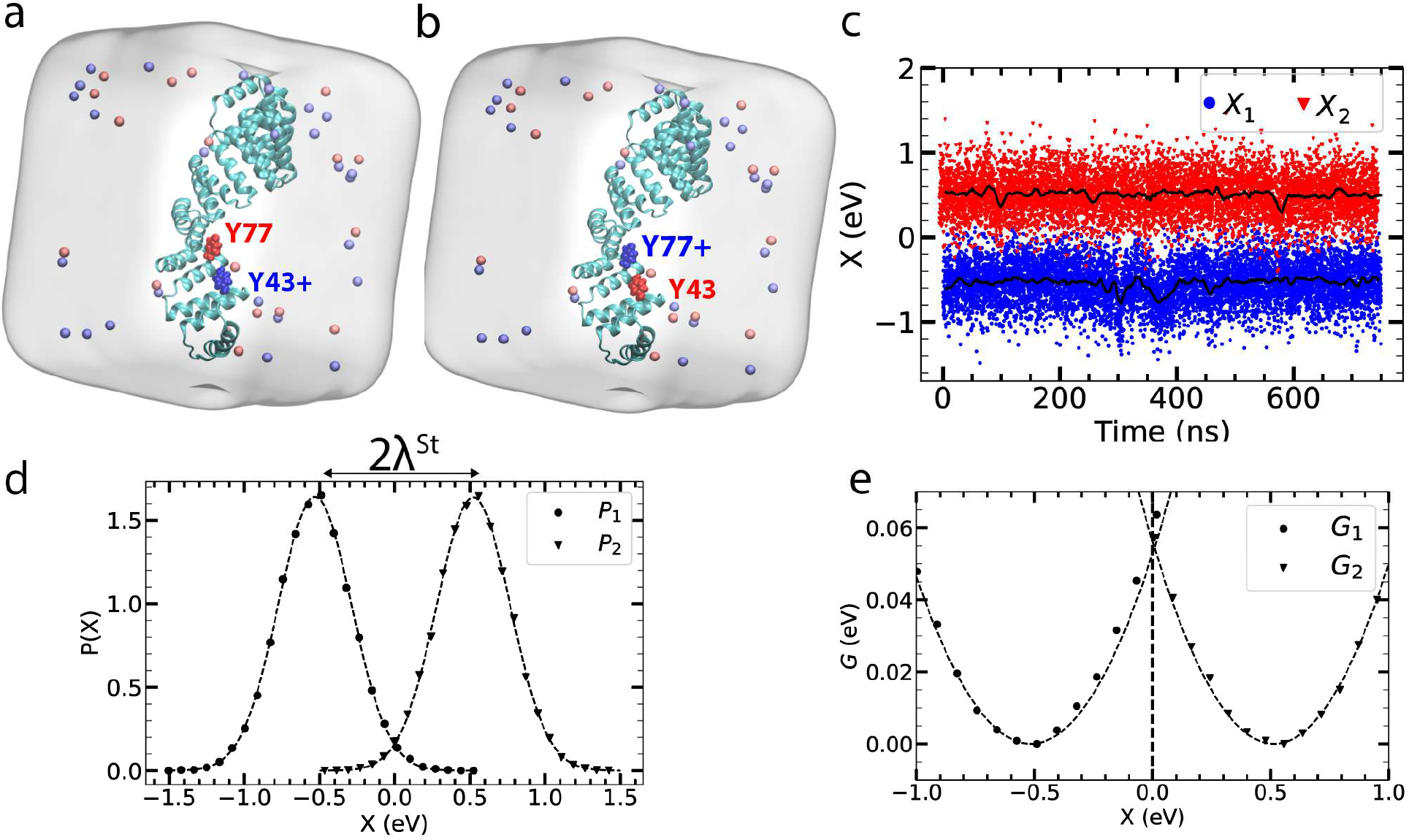
MD simulations of the electrostatic energy gap produced by a charge transfer process: **a)** All-atom simulation system representing the initial state of the Y77-to-Y43 charge transfer process, with Y43 being positively charged (oxidized) and Y77 being electrically neutral. **b)** Final state of the Y77-to-Y43 transfer, with Y43 being neutral and Y77 positively charged. **c**) Energy gap fluctuations at Y43 (X1) and Y77 (X2). The black traces indicate 10-ns block average of the 10-ps sampled raw data. (**d**) Electrostatic energy probability distributions for the positively charged Y43 (triangles) and positively charged Y77 (circles). The separation between the distributions’ peaks is 2λ^St^. **e)** Free energy as a function of the donor-acceptor electrostatic energy gap: the intersection at X=0 defines the transition state.

Electron transfer occurs at the point of crossing of these two distributions (*X*=0 in Figure 3e). The variance of the distribution yields λ from 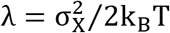. For this particular D-A pair, the full width of the *X* distribution is ~ 0.5 eV yielding *σ*~eV and λ~1.15 eV. Thus, the free-energy parabolas are wider than the Stokes shift (λ^St^~ 0.52 eV Figure 3d) would predict. For this particular transfer reaction, Eq. 2 yields *λ*^*r*^ = 0.24 eV, substantially less that the value derived either from the Stokes shift or from the distribution variance.

Similar calculations (see Methods) were carried out for all pairs of aromatic residues within the Dutton radius (1.4 nm edge to edge distance in the crystal structure) within two repeat units, which, together with the periodicity of the CTRP8 protein, encompassed all likely electron transfer paths. Test simulations carried out for equivalent pairs of residues from different protein units have explicitly verified the convergence of our rate calculations, Figure S1. Thus, taking the repeat structure of the protein into account, we only needed to calculate the activation energies for residue pairs within one protein unit and between the two adjacent units.

Figure 4a and b show cumulative averages of the MD-determined reorganization energy values for a representative Y-Y and W-Y residue pair. Note here that λ1 (orange trace) is for the D-A^+^ state and λ2 (green trace) is for the D^+^-A state (defined in Figure 2c). In either configuration, λ is the solvation free energy of the electron transfer dipole, which is “−“ on the acceptor and “+” on the donor. Since the charge distribution is the same, λ is expected to be the same in both states and indeed *λ*_1_ ≈ *λ*_2_. The average of the two values is used in what follows.

**Figure 4:**
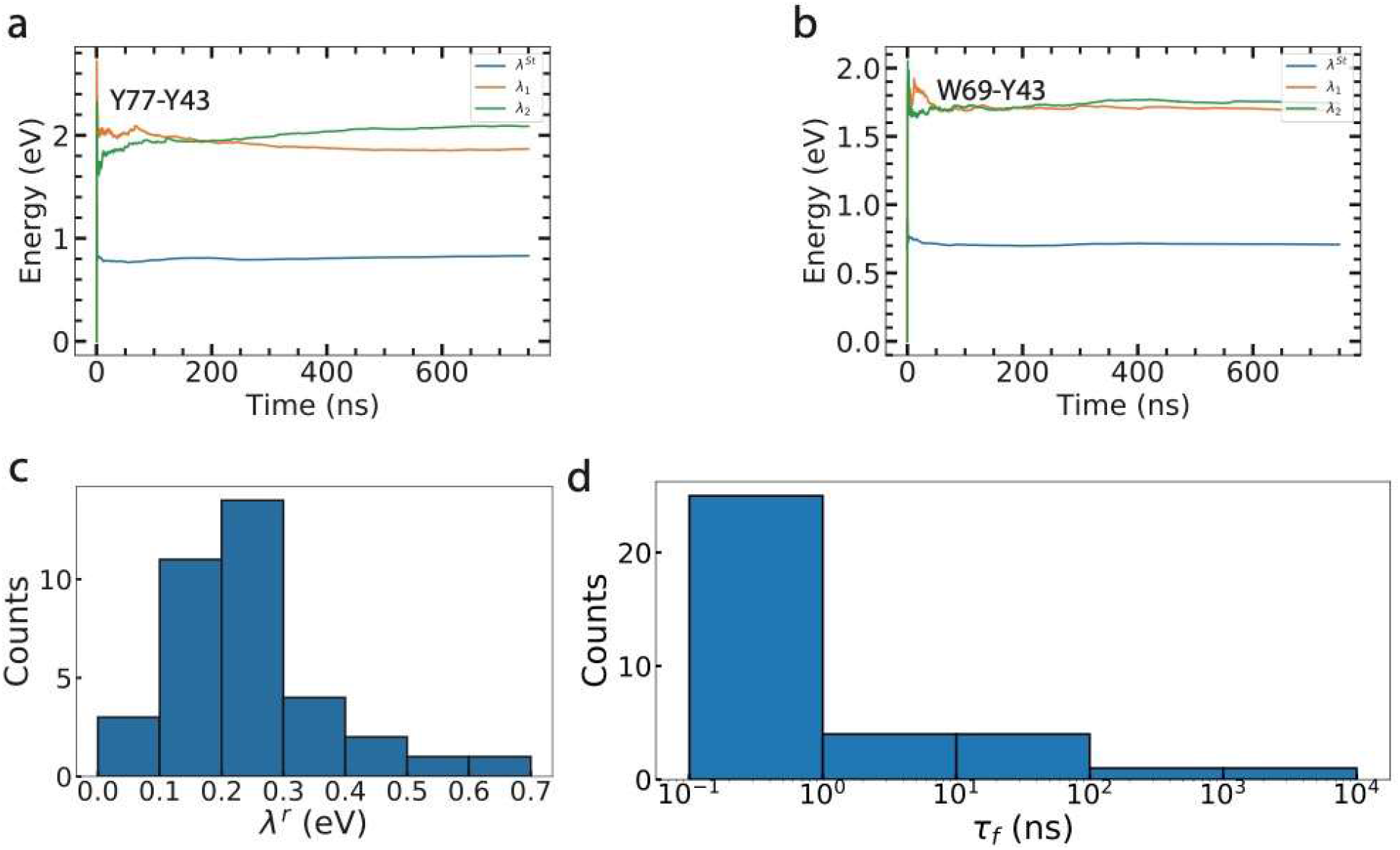
Calculation of reorganization energies: **a,b)** Reorganization energies for a tyrosine-tyrosine (panel a) and a tyrosine-tryptophan (panel b) electron transfer calculated as 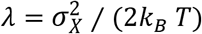. The yellow and orange curves show cumulative average values of λ for the two charge shift reactions. The two values should be equal, in principle, and hence their average,*λ*, is used in subsequent calculations. The blue curve indicates *λ*^*St*^. As *λ* is substantially greater than *λ*^*St*^, *λ*^*r*^ = (*λ*^*St*^)2/*λ* is substantially smaller than *λ*^*St*^. **c)** Distribution of *λ*^*r*^ values calculated for all pairs of aromatic residues located within the Dutton radius of each other. The modal value of the distribution is 0.2 eV, four times smaller than 0.8 eV derived from the Stokes shift. **d)** Distribution of the hopping times calculated from the distributions of *λ*^*r*^, the corresponding edge-to-edge donor-acceptor distances and using the polarizability adjusted reorganization energies. The modal time is ~ 100 ps, consistent with the experimentally observed nA currents.

The distribution of *λ*^*r*^values, computed according to Eq. 2, is plotted in Figure 4c. The modal number extracted from our simulations was 0.2 eV. A list of the values of *λ*^st^, *λ*_*var*_, Δ*G*, *λ*^*r*^ and edge-to edge distances for all interacting pairs within a Dutton radius is given in Table S1. As these simulations employed a nonpolarizable force field, they neglected possible contributions of induced dipoles that would screen the electrostatic interactions, leading to even further reduction of the reorganization energy. To account for such electronic screening, we introduced a correction factor of 0.8, as discussed in detail in Supplementary Information. The correction factor was applied to both *λ*^St^ and *λ*, yielding an even lower modal value of *λ*^*r*^ of 0.16 eV, substantially smaller than 0.8 eV, a value commonly accepted for proteins. The activation barrier (for ΔG = 0) is *λ*^*r*^/4, so *λ*^*r*^ = 0.16 eV gives an activation barrier of 40 meV, close to thermal energy at room temperature. Overall, these data yield an average value for 〈*λ*^*St*^/*λ*〉 ≈ 0.4. Since the oxidation potential of tyrosine and tryptophan in solution includes a reorganization energy between 0.5 and 2 eV (17, 18), the reduction of reorganization energy in a protein-aqueous medium to 0.16 eV could account for the resonant injection of charge (Figure 1e and Eq. 3) because the potential of the (non-equilibrium) oxidized state will be shifted towards the vacuum by ~ 0.7 eV.

Our calculation of the hopping rates proceeds as follows. The charge transfer rate *k*_*ET*_ is given by

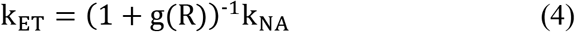

where the non-adiabatic transfer rate,*k*_*NA*_, is given by

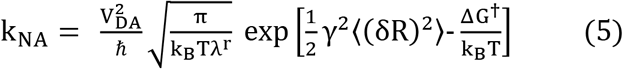

and the activation barrier ΔG^†^is given by Eq. 1 in which *λ* is replaced with *λ*^*r*^ and the reaction free energy is from Eq. 3. 〈(*δR*)^2^〉 in is the variance of the donor-acceptor center of mass distance. The crossover parameter *g*(*R*) (see Methods and Supplementary Information, Eqs. S1-S7) accounts for solvent dynamical control (34) of the rate pre-exponential factor (35–37). The factor *g*(*R*) includes two dynamic components: the Stokes shift of the energy gap *X*(*t*) and the donor-acceptor distance, *R*(*t*), modulating the donor-acceptor electronic coupling *V*_*DA*_. *V*_DA_ decays exponentially with the donor-acceptor edge to edge distance and is calculated here from the Hopfield equation (38)

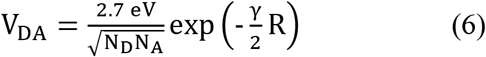

where *R* is the edge-to-edge distance (in nm) and *N*_*D,A*_ =7 for a tyrosine and 9 for a tryptophan. *γ*/2 is the distance decay parameter, equal to 7.7 *nm*^−1^ (38). Taken together with the activation barriers obtained from the MD simulations, these equations yield the hopping times for all pairs within the Dutton radius in the protein, and the distribution of these times is plotted in Figure 4d and the forward and backward rates are listed in Table S2. Although the overall electric current is determined by the carrier velocities, and not by the transit times *per se*, the hopping times of ~0.1 ns (Figure 4d) are consistent with the nA currents measured in these molecules and are much smaller than 1μs estimated for λ = 0.8 eV. The dynamical crossover parameter 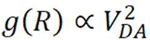 gains its distance dependence from the electronic coupling. Therefore, for *g*(*R*) > 1, the term 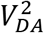 cancels in Eq. 4 and the rate constant becomes insensitive to the electronic coupling. Many electron hops fall in this regime of dynamical control (Table S2): for instance, charge transfer between Y77 and Y43 shown in Figure 3 produces *g* ≈ 28. The dynamical control introduces the dependence of the electron-transfer kinetics on protein dynamics and elasticity which are absent in traditional theories.

### Calculation of currents and current decay rate from the hopping rates

The decay of current with distance is described by a parameter that depends on the electronic coupling between the electrodes and the entry/exit residues, the size of those residues, and the diffusion constant, as we will show below. The electronic coupling can be estimated from the measured contact resistance, and the size of the entry/exit residues is known, leaving the diffusion constant to be determined.

A model of the electrode-protein system is shown in Figure 5a, from which we derive an expression for the current through the complex, below. Electrons are extracted from the residue closest to the right (positive) electrode (2 on Figure 5a) and injected in a similar way from the residue closest to the left (negative) electrode (1 in Figure 5a). The experimentally determined contact resistance (Figure 1d) is a significant fraction of the overall resistance, particularly for the shorter molecules. The fraction, *K*(*L*), of the applied bias dropped across the bulk of the molecule was determined using an elementary circuit analysis, (13) and a fit to *K*(*L*) is shown in Figure S2. However, even in the longest molecules, where a significant fraction of the bias is dropped across the protein, the applied field is small compared to the internal fields within the protein (see, for example, Figure 4 of Martin et al. (39)). Thus, once injected, the charges diffuse under the driving force of the carrier gradient owing to electron injection at contact 1, until captured by contact 2, a process characterized by the carrier diffusion constant.

**Figure 5:**
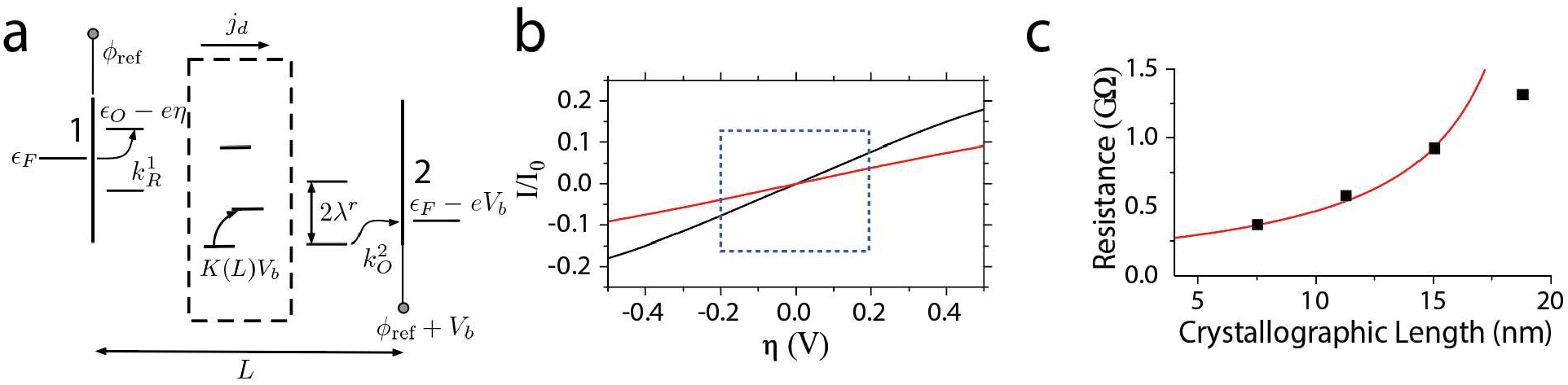
Charge injection model of the current: **a)** An electron is extracted at the right contact 2 (energy *ϵ_F_* − *eV_b_*) at a rate 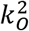 via the oxidation of the closest aromatic residue. The oxidized state then diffuses, through a sequence of hops, within the body of the protein (dashed box, current density *j*_*d*_) to be collected by the contact 1 on the left via reduction of the closest residue at a rate 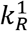. *ϵ_F_* is the Fermi energy of the electrodes, *V_b_* the applied bias, *η* the potential drop at the contacts and *ϕ*_*ref*_ the potential of the electrodes with respect to a reference electrode. The arrows show the direction of electron motion. *K(L)* is the experimentally-determined fraction of the bias that is dropped across the bulk of the protein (Figure S2). **b)** Calculated current-voltage dependences for L=10 nm (black) and L=15 nm (red) where the current is expressed in units of the saturation current I_0_. The curves are linear in the ±200mV range (blue box), as observed in the experiments (Figure 1c). **c)** Calculated resistance vs length (red line) fits experimental data (dots) with I_0_ = 5.5 nA, and the parameter *a* = 3.8 (see text for detail). the combination of the distance dependence of hopping diffusional current with the length-dependent potential drop across the contacts.

### Diffusion constant

The diffusion constant connects the diffusion flux to the gradient of the volume density of charge carriers in the macroscopic Fick’s law. A one-dimensional Brownian walker stepping the distance Δ*x* in each step requiring time *τ* has the diffusion constant *D* = (Δ*x*)^2^/(2*τ*). The challenge of extending this equation to carrier diffusion in the protein is that both the step Δ*x* and the hop time *τ* become distributed variables. The problem is solved analytically by Derrida’s model (40) which sets up diffusion on a one-dimensional periodically replicated chain of sites with fixed Δ*x* and forward and backward transition rates specified at each lattice site. This solution does not allow distributed values of Δ*x* and provides no algorithm for choosing a path maximizing diffusion. The latter problem is resolved by kinetic Monte Carlo (KMC), which generates multiple paths through the network of Tyr and Trp residues and yields the time of first passage between the injection and collection sites. However, it does not provide a direct formalism for calculating the diffusion coefficient. The ability of the carriers to tranverse the 3D protein over many alternative paths is an essential aspect of the problem that has to be captured by a successful formalism. In the absence of such a formulation, we adopted a somewhat heuristic extension of the result for a Brownian walker (to which Derrida’s model reduces for symmetric forward and backward rates), calculating the diffusion constant in the form

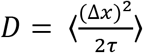

where both the step size Δ*x* and the step time *τ* are stochastic variables changing along the paths produced by KMC. Specifically, *τ* is calculated as the waiting time on a given site. In the limit of a single path, this result reduces to Deridda’s calculation, where the average is now for the sites along the path. In the KMC formulation, the average was taken along a given path, with the most probable value of diffusion constant given by the peak of the distribution calculated for all paths.

The calculation is set up using the hopping rates and center-of-mass distances (Table S2) for all aromatic pairs within a Dutton radius in two repeats of the CTPR unit. We use two repeats so as to include the hops between neighboring repeats (c.f., Figure 2b). As shown in Figure S1, the properties of longer proteins are captured via translational symmetry.

Using the data listed in Table S2, a graph was constructed with edges that represent the hopping rates, with two edges between each pair of residues, one representing the forward hopping rate and the second the backward hopping rate. Electron hopping MC simulations were run until the electron reached the exit residue, with the value of *D*_*i*_ calculated at the *i*^th^ hop from the residence time, *τ*_*i*_, and the center of mass hopping distance Δ*x*_*i*_. On completion of the passage, an average of these values is stored (Eqns. S17-21).

In the experiments, N- and C-terminal cysteines were incorporated into the proteins in order to form defined chemical contacts to the electrodes. Examination of the structure (Figure 2b) shows that C-terminal injection or extraction is only possible at Y89. Injection at the N-terminal can occur at W35 (the closest residue in sequence to the N terminus) or Y36 which is at almost the same through-space edge to edge distance (0.9 nm) from the cysteine (W35 is 1 nm through-space edge to edge distance from the cysteine). We have therefore calculated the distribution of *D* values for paths that start or end on either of these residues (the inclusion of backward and forward hops captures diffusion in either direction). 100000 simulations were run for each set of paths and the distributions of *D* are shown in Figures 6a (W35 path) and 6b (Y36 path). The corresponding most probable *D* values are 22.1 nm^2^/ns and 22.8 nm^2^/ns.

**Figure 6:**
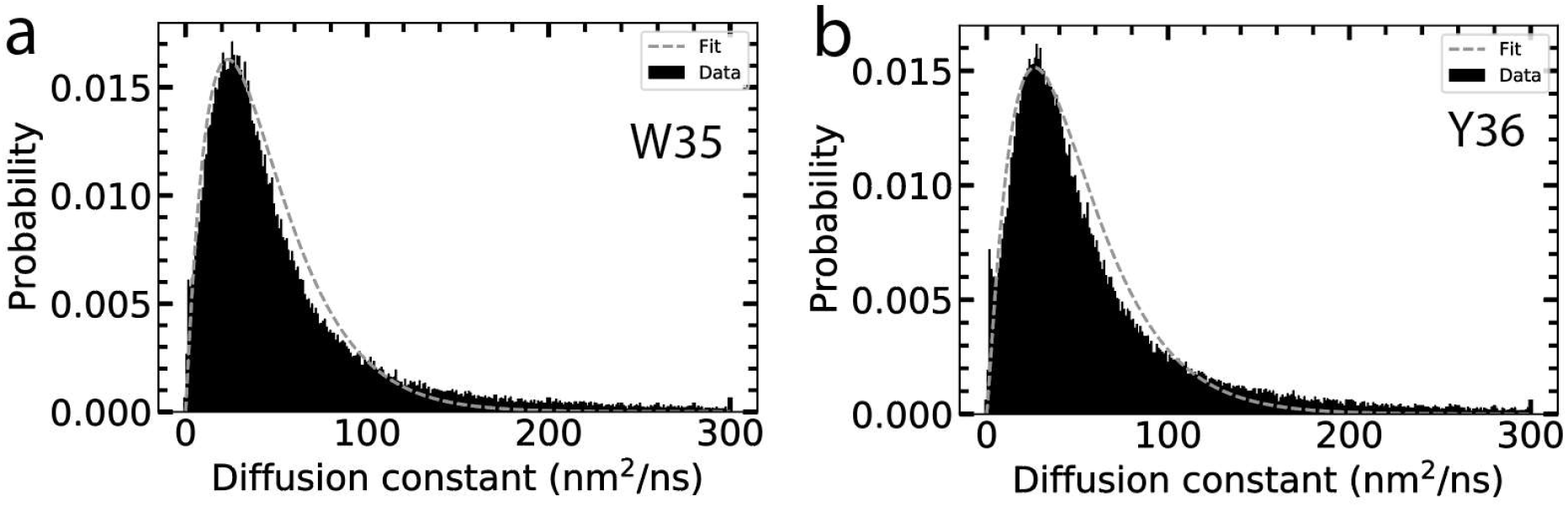
Distribution of diffusion constant values for paths that connect W35 and Y89 (a) and Y36 and Y89 (b) calculated via a kinetic Monte Carlo algorithm. Fitting with Gamma function distributions (dashed lines) yields most probable values of *D* = 22.1 nm^2^/ns for the W35 paths and 22.8 nm^2^/ns for the Y36 paths.

As a check on this heuristic approach, we used the Derrida formalism (40) to calculate values for two limiting cases. In one case we chose a single path that maximized forward rates (which is not the same as maximizing the diffusion constant) and calculated the corresponding diffusion constant (this path illustrated in Figure 2b). The calculation was carried out following Eqns. S13 and S14 using the rates shown in Table S3 (the final hop that closes the loop as required for the Derrida theory is Y89 to W35, equivalent to Y89 to W103 owing to translational symmetry – Figure S1). For this path we obtain *D* = 3.06 nm^2^/ns, an order of magnitude less than the multiple-path value obtained from the KMC calculations. An alternative approach, using the Derrida theory, is to find all the paths that connect entry and exit, calculate the corresponding *D* values for each, and summing them, on the assumption that each path adds a current proportional to its corresponding *D*. The obvious fault with this approach is that paths are allowed to overlap – i.e., this approach allows a given site to be multiply oxidized (which is energetically impossible). This will yield an overestimate of *D*. We find 433 such paths connecting Y36 to Y89 for which the total *D* = 578 nm^2^/ns, an order of magnitude larger than the value obtained from the MC calculations. These two limiting cases bound the results of the MC simulations, indicating that our heuristic calculation of *D* from KMC is likely of the correct order of magnitude.

#### Charge injection

We assume charge injection into a single site for which the injection and recombination rates depend only on the potential drop at the contacts, *η* (Figure 5a) and the electronic coupling between the electrodes and the entry and exit sites, assumed, for simplicty, to be the same, Δ eV, at each contact. Using empirical data to calculate *η* (Figure S2) and expressions for the injection rates (Eqns. S15) solved for stationary conditions (Eqns. S14-16) results in the following expression for the contact conductance:

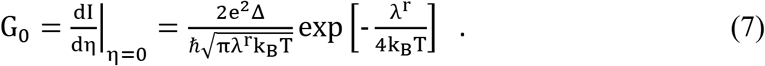

Evaluating this expression for *λ*^*r*^ = 0.16 eV and a measured contact conductance of 4.55 nS (1/220 MΩ) yields Δ = 5 × 10^−6^ eV.

#### Stationary current

The stationary current through the protein can be written in terms of the fraction of oxidized states at each contact *n*_*i*_ and the diffusion flux *J*_*d*_ of charge carriers through the protein with *n*_*i*_ satisfying the following current balance conditions:

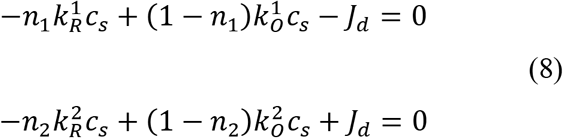

where J_*d*_ = −D ∂_*x*_ρ(x) is the diffusional flux due to the gradient of the bulk number density *ρ(x)* and *D* is the diffusion constant calculated above. Expressing the rates 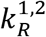 and 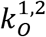 (Eqns. S15) in dimensionless form leads to

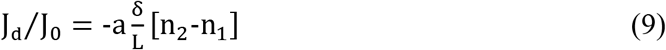

where *J*_0_ = c_*s*_ Δ/ℏ = Δ/(ℏ*S*) and *S* is the contact area. The saturation current is given by I_0_ = eJ_0_S = eΔ/ℏ ≈1.2 nA (using Δ = 5 × 10^−6^ eV). The dimensionless parameter *a* is given by

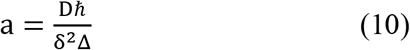

Here, *δ* is the size of the entry/exit residue (adopted as 0.96 nm, representative of the two residues). With the values of *D* calculated above (22.1-22.8 nm^2^/ns), we obtain *a* = 3.12 – 3.22.

Eqns. 8 and 9 can be solved for and to produce the diffusive current of holes and the corresponding conductivity:

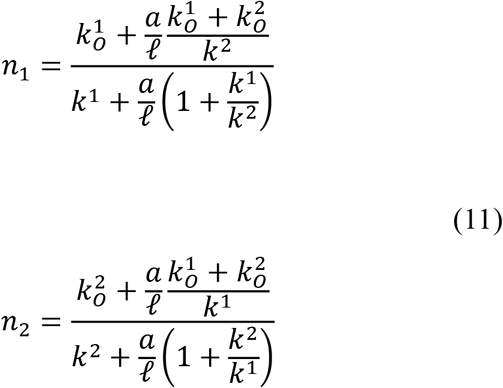

where k^1^ = k_*O*_(η) + k_R_(η) and k^2^ = k_O_(−η) + k_R_(−η) as calculated from Eqns. S14, and ℓ = *L*/*δ*.

Equations 11 provide the solution for the stationary current through the protein:

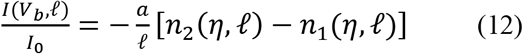

where the dependence of *η* on *V*_b_ and ℓ is given in the caption to Figure S2. Figure 5b plots 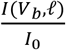 vs *V*_b_ for two protein lengths (10 and 15 nm). With *I*_0_ ≈ 1.2 nA (as calculated from the contact resistance above) the calculated I-V curves are of the same magnitude as the examples given in Figure 1c, and feature a linear regime in the ± 200mV range as observed experimentally.

The resistance as a function of ℓ is given by

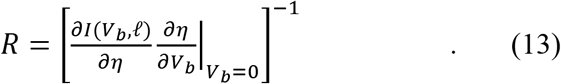

This function is fitted (red line) to the experimental data (squares) in Figure 5c, with the best fit parameter *a* = 3.8, close to the values (3.12 – 3.22) calculated above, showing that the present approach correctly predicts the current decay with distance. In a previous experimental work (13), the non-linear dependence of the molecular resistance on length was hypothesized to be intrinsic to the difussion process. The analysis presented here predicts an inherently linear dependence, as it must be, given that the current is proportional to the carrier velocity, i.e., the ratio of diffusion time to molecular length. The observed non-linearity is mostly accounted for by the dependence of the potential distribution on the length of the molecule.

The absolute magnitude of the current is determined by *I*_0_ which was calculated to be 1.2 nA via the estimate of the electronic coupling constant, Δ, a parameter that the decay rate (Eqn. 10) also depends on. This estimate is based on the assumption of equal resistance of two contacts which can be merged into one residue to calculate the contact resistance. The fit to the resistance data (which are based on the peak of the distributions obtained from many current-voltage curves) is based instead on evaluating the diffusive current in Eqs. 9 and 12 and required *I*_0_ = 5.5 nA, ~ 4× the value obtained from Δ, but surprisingly close, given the limitations of the contact model.

The rather precise agreement between the calculated decay rate *a* = 3.12 – 3.22, and the value obtained from experimental data (a= 3.8) is probably fortuitous, given both the lack of an exact theory for the difussion process and other approximations used here: The force fields employed in MD are nonpolartizable and thus neglect screening of charges by induced molecular dipoles. An indication that this can cause minor inconsistencies is the fact that the mean ratio 〈*λ*^St^/*λ*〉 ≈ 0.4 is somewhat higher than ≈ 0.3 from comparing Eq. 3 with the position of the conductance peak in Figure 1e. Dynamical hetergeneity, in particular at sites closer to the protein-water interface, can lead to faster dynamics and, correspondingly, to lower *g*(*R*) in Eq. 4. Despite these limitations, the present theory demonstrates how long-range hopping is enabled through a reduced reorganization energy owing to non-ergodicity. The theory predicts the correct magnitude of current, and accurately predicts its decay with the length of the proteins.

## Conclusions

The key result of this work is that proteins that do not contain redox cofactors can support long-range and rapid hopping of electrons, with implications for the role of aromatic amino acids in facilitating charge transport. The same reduction of reorganiziation energy that accounts for this long-range transport also accounts for the alignment of the non-equilibrium charge shift states with the Fermi energy of some noble metals, important for the use of proteins as bioelectronic components. The reorganization energy appropriate to non-equilibrium oxidation states of tyrosine and tryptophan, ~ 0.16 eV, is significantly smaller than the Marcus reorganization energy derived from the Stokes shift (~0.8 eV). This reduced reorganization energy corresponds to a barrier (for of 40 meV, which is close to available thermal energy. This reduction of reorganization energy has previously been demonstrated for charge transfer between cofactors (22, 24, 25) and is now extended to the aromatic amino acids. Even in the case of proteins rich in cofactors, the origin of the required reduction in reorganization energy remains obscure (15), so it is interesting to note that a reorganization energy below 0.2 eV and a fast diffusion constant (> 20 nm^2^/ns), the conditions required for fast transport in OmcS proteins, are obtained in the present case for transport a protein that lacks any redox cofactors. Finally, this reduction of the reorganization energy provides a plausible explanation for the fact that the current carrying states appear to be some 700 mV closer to the vacuum (i.e., at 300 mV vs NHE) than would be expected from the equilibrium oxidation potentials of tyrosine and tryptophan (1V vs NHE).

## Methods

### MD protocols

All MD simulations were performed using NAMD2 (41) periodic boundary conditions, the CHARMM36 force field (42) for the protein and the TIP3P model (43) for water. Van der Waals and short-range electrostatic forces were evaluated using a 1.2 nm cutoff whereas long range electrostatic forces were evaluated using the particle mesh Ewald method computed over a 0.12 nm-spaced grid (32). The initial equilibration of the CTPR8 system was performed using a 2 fs time step. All simulations of the CTPR8 systems containing an oxidized residue were performed using a 4 fs integration time step and the hydrogen mass repartition scheme (33). The equilibration simulations were performed in the constant number of particles, pressure (1 bar), and temperature (310K) ensemble maintained using a Langevin dynamics thermostat and the Nose-Hoover Langevin pressure control (44, 45). Visualization and analysis were performed using VMD (32).

### MD simulations of CTPR8 systems

The atomic coordinates of a CTPR8 protein were taken from the published crystal structure (46). To reproduce the protein construct used in experiment (13), the N and C termini of the protein were extended by 31 and 18 amino acids, respectively, both terminating with a cysteine residue. The partially folded structure of the N-terminal addition was generated using the Iterative Threading ASSEmbly Refinement (I-TASSER) server (Zhang lab, University of Michigan) and its default parameters (47), the C-terminal addition was assumed to be disordered. The resulting atomic model of the experimental CTPR8 protein is provided as supplementary material, CTPR8-Cys.pdb. Missing hydrogen atoms were added to the protein using the psfgen plugin of VMD. The protein was immersed in a 8.0 x 8.6 x 7.1 nm^3^ pre-equilibrated volume of water using the Solvate Plugin of VMD (48). Potassium and chloride ions were added using the Autoionize plugin to produce 150 mM KCl solution. The system was minimized using the conjugate gradient method for 100 steps, which was followed by a 500 ns equilibration run. During the first 25 ns of the equilibration, the Cα atoms of the protein were restrained to their initial coordinates using harmonic potentials with the force constant of 100 kcal/(mol nm^2^). The microscopic configuration of CTPR8 system obtained at the end of the 500 ns equilibration was used to construct thirteen CTPR8 systems each containing a different tryptophan or tyrosine residue in its oxidized state. These tyrosine and tryptophan residues were assumed to be oxidized to their cationic free radical state, carrying an overall charge of one proton. The partial charges of an oxidized tyrosine or an oxidized tryptophan residue were taken previous semi-empirical (29) and quantum mechanics/molecular mechanics (QM/MM) (30) calculations, respectively. The patches used to oxidize the residues are provided as supplementary material (oxidize_Y.txt and oxidize_W.txt), the values of the partial charges are listed in Table S4 and S5. Each system containing an oxidized residue was then equilibrated for 750 ns. Atomic coordinates were saved every 10 ps.

### Calculation of the electrostatic potential at the active residue

To calculate the energy-gap trajectory, we evaluated the electrostatic potentials at the location of the atoms of the redox-active residue every 10 ps using the method (32) implemented in the PMEpot Plugin of VMD. Briefly, every partial charge in the system was approximated by a spherical Gaussian

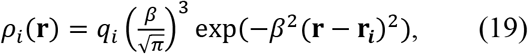

where parameter *β* is the width of the Gaussian. The instantaneous electrostatic potential corresponding to the charge density was obtained by solving the Poisson equation

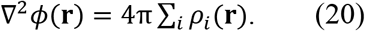

The resulting electrostatic potentials Φ_*j*_ at the location of atoms *j* (atoms whose charge changes during the charge transition) were extracted from the MD simulation of the oxidized and reduced residues. The electrostatic (Coulomb) component of the donor-acceptor energy gap was calculated as

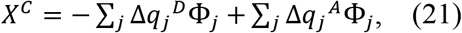

where 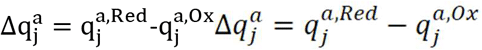 are differences of atomic charges in the reduced and oxidized states of the donor, a=D, and acceptor, a=A, moieties.

The Stokes-shift reorganization energy is given by the difference of average values of the energy gap in the initial and final charge-transfer states

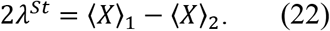

The average 〈*X*〉_1_ was calculated from MD simulation of the reduced donor and oxidized acceptor whereas 〈*X*〉_2_ was calculated from MD simulation of the oxidized donor and oxidized acceptor (see Figure 2c for the reaction diagram). The variance reorganization energy *λ* was determined as the mean of the energy-gap variances 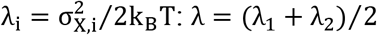. The hopping rates (Table S2) were calculated according to Eqns. 4-5.

### Evaluation of the dynamic crossover parameter g

The parameter g is given by (34)

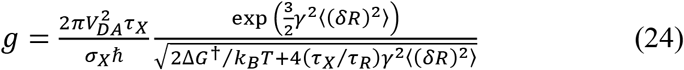

where ΔG^†^ is given by Eq. 1 in which *λ* is replaced with *λ*^*r*^ and the reaction free energy is from Eq. 3, 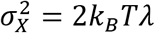, and *τ_X_* and *τ_R_* are the relaxation times of the Stokes-shift dynamics and of the donor-acceptor distance, respectively. The integrated relaxation times *τ_X,R_* were calculated using the time correlation functions from MD (Figs S3 and S3 and Eqns. S6 and S7).

## Acknowledgements

We thank Eathen Ryan for generating the protein structures. This work was supported by grant 1R01HG011079 from the National Human Genome Research Institute, grant CHE-2154465 from the National Science Foundation, grant W911NF2010320 from the US Army and the Edward and Nadine Carson Endowment. AA acknowledges support via grants from the National Science Foundation (PHY-1430124 and DMR-1827346). The supercomputer time was provided through the Extreme Science and Engineering Discovery Environment (XSEDE) allocation MCA05S028.

## Supplementary Information

**Figure S1.**
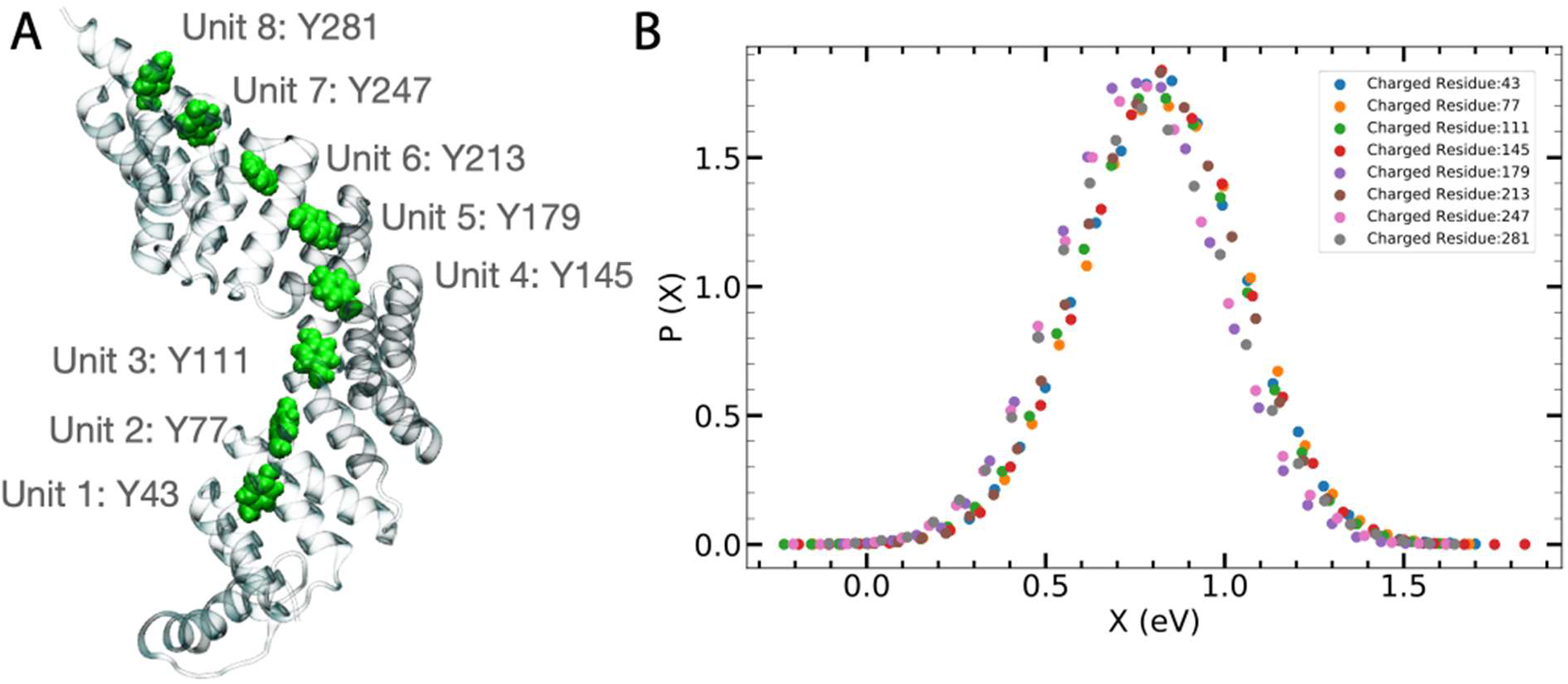
Energy gap fluctuations of equivalent residues in CTPR. **A)** Y43 and its equivalent residues in the eight repeat units of CTPR8. The sidechain of the equivalent residues are highlighted using green spheres. The unit number for each of the residues is indicated next to the residue. **B)** Electrostatic energy gap fluctuation for the oxidation half reaction computed for equivalent tyrosine residues from the eight repeat subunits of CTPR. The distributions have been normalized.

**Figure S2:**
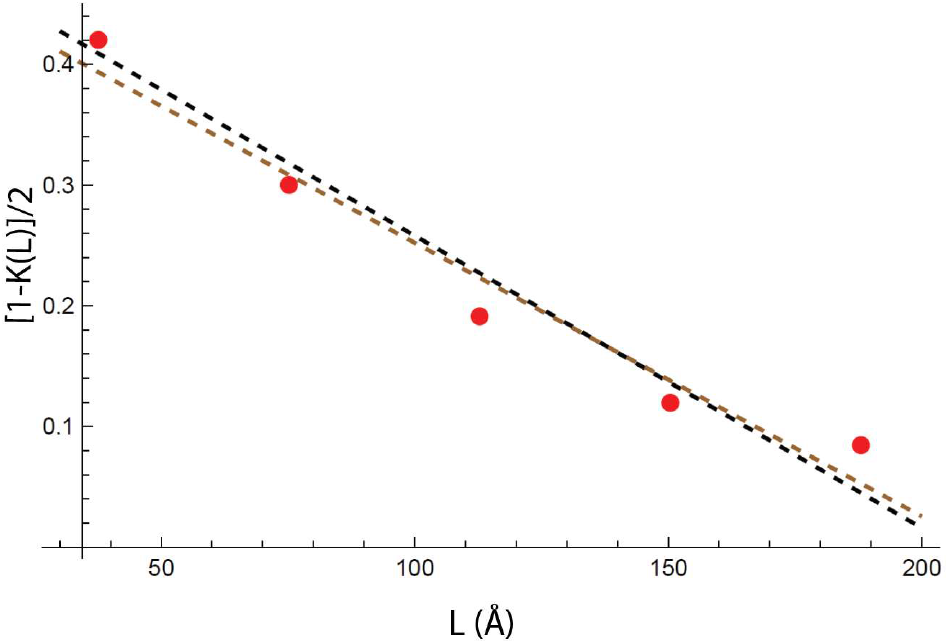
Linear fit to the experimentally-determined bias distribution. K(L) is the fraction of the applied bias that appears across the molecule. The potential drop at the contacts is 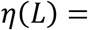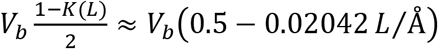.

### Stokes-shift and Distance Dynamics

The dynamical crossover parameter *g* accounts for the solvent dynamical control of the rate pre-exponential factor. Equation S1 shows that *g* includes two relaxation times: the integral relaxation times τX for the Stokes-shift dynamics and the relaxation time τR for the dynamics of the do-nor-acceptor distance^7^

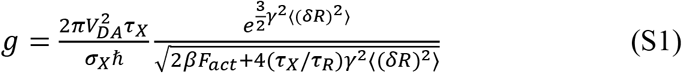

In addition,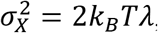, γ is the parameter of the exponential fall-off of the squared donor-acceptor electronic coupling and 〈(*δR*)^2^〉 is the distance variance.

The Hopfield equation^8^ used to calculate *V*_DA_ is based on the edge-to-edge distance (*R*_edge_, in nm) between the donor and acceptor

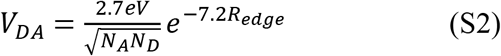

Here *N*_D,A_ = 7 for tyrosine and *N*_D,A_ = 9 for tryptophan. In addition, the activation energy ΔG^†^ in Eq. 1 is calculated as

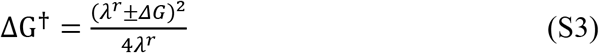

with “+’’ and “−” referring to the forward and backward transitions. In Eq. S3, *λ*^*r*^ is the reaction (superscript “r”) reorganization energy calculated from the equation^9^

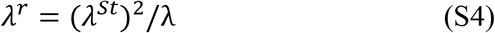

where the variance reorganization energy 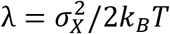 is calculated from the variance of the energy gap reaction coordinate *X*. We take the mean of the forward and backward *λ* to calculate the reaction reorganization energy *λ*^*r*^ in Eq. S4. The Stokes-shift reorganization energy *λ*^St^ is given by the difference of average values of the energy gap in the initial and final electron-transfer states

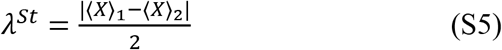

Calculations of the Stokes-shift and donor-acceptor distance dynamics were based on MD simulations using NAMD^2^ with a 4 fs integration time step and with hydrogen mass repartition.^10^ Simulations in the NPT ensemble were performed using a Langevin dynamics thermostat and the Nose-Hoover Langevin^4,5^ piston pressure control set at 310 K and 1 atm. The Stokes-shift dynamics were calculated using two MD trajectories, one being 1 ns in duration and sampled every 12 fs and another being 250 ns in duration and sampled every 10 ps. Both simulation systems probed the same electron transfer reaction between oxidized residue Y43 and neutral Y77. The energy gap was calculated as a function of time for the short and long trajectories. The two trajectories were combined while disposing the first 1ns of the longer trajectory. The Stokes-shift correlation function *C*_*X*_(*t*)= <δX(t)δX(0)> was normalized and fitted to a sum of a Gaussian ballistic decay and two exponential functions

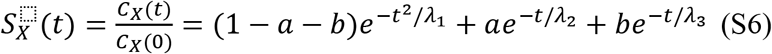

The MD data and the fit are shown in Fig S3. The fit was obtained with the parameters: a = 0.085, b = 0.662, *λ*_1_= 0.0777 ns^2^, *λ*_2_ = 3.8 × 10^−3^ ns, and *λ*_3_= 6 × 10^−5^ ns. These fitting parameters give the integral relaxation time of 20 ps.

The relaxation time for the dynamics of the donor-acceptor distance was calculated from a 65 ns trajectory. The center-of mass distance R(t) between Y55—Y76 residue pair was sampled with the time step of 2.5 ps producing the correlation function

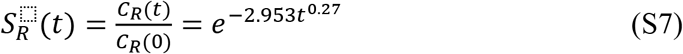

where *C*_R_(*t*)= <δR(t)δR(0)>. The result is shown in Fig S4. The fit to Eq. S7 yields the integral relaxation time *τ*_R_ = 281 ps.

**Figure S3:**
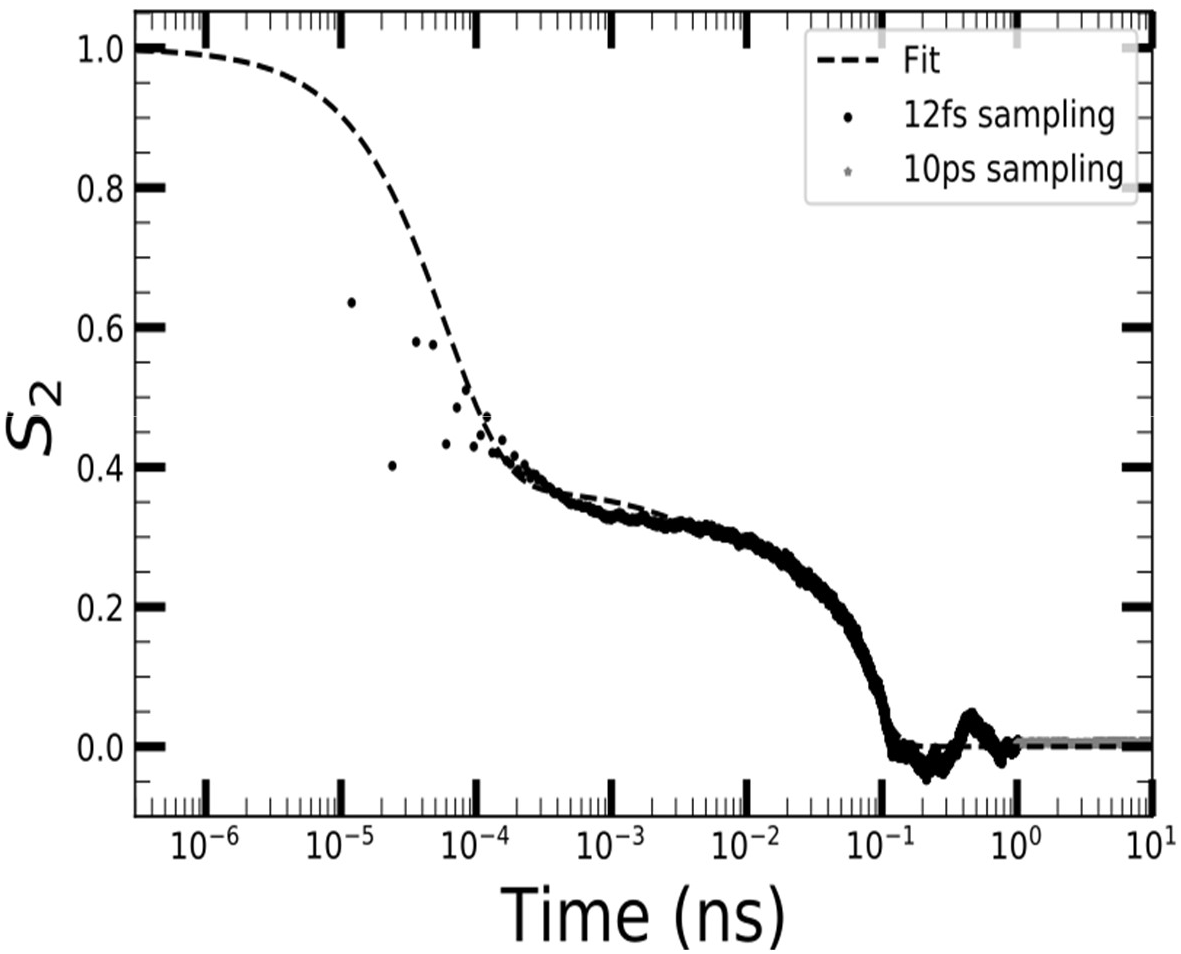
Stokes-shift time correlation function. The dots and stars are the data from MD simulation, while the dashed line indicates the fit to Eq. 6. The grey stars indicate correlation data from 10ps sampling step and the black dots indicate correlation data during the 12fs step.

**Figure S4:**
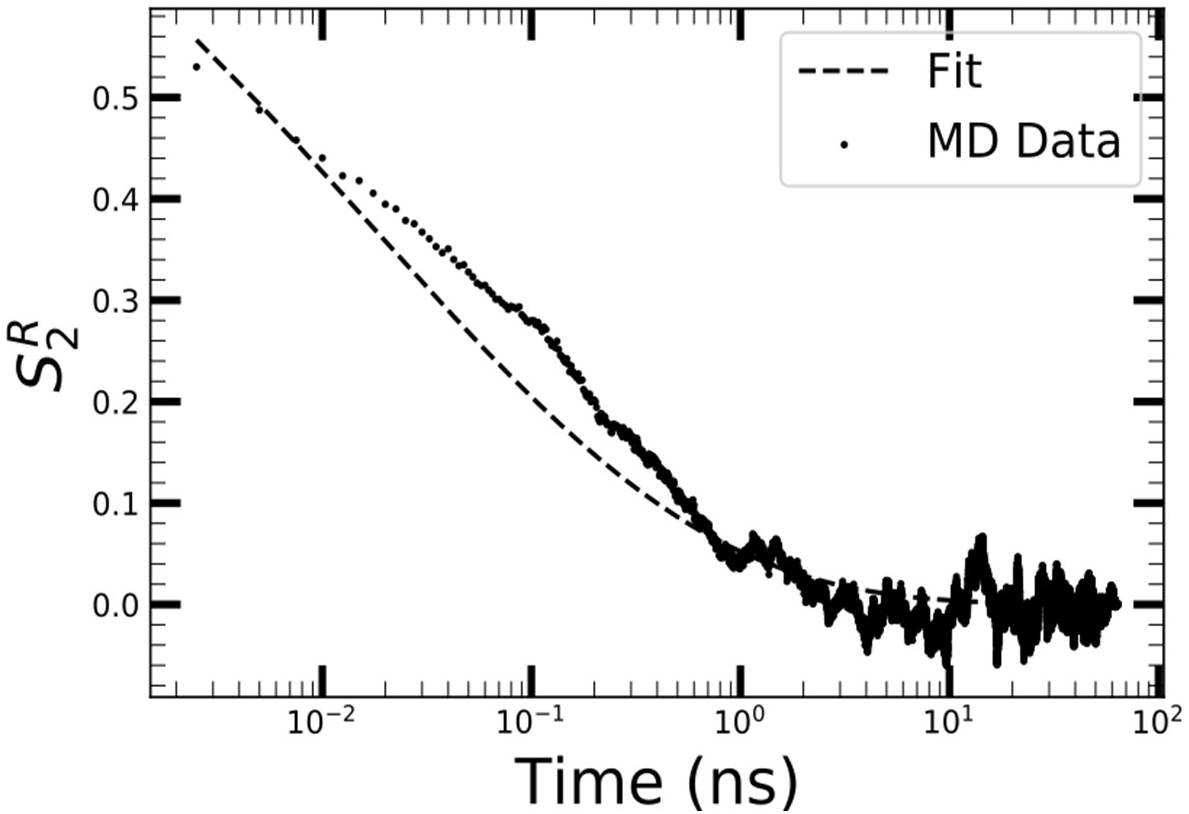
Distance time correlation function. The dots are the data from MD simulation, while the dashed line indicates the fit to Eq. 7.

### Polarizability

We conduct the MD simulations in non-polarizable force fields, which do not account for screening of electrostatic interaction by electronic induced dipole moments. The standard dielectric theories account for the medium polarization in terms of the Pekar factor 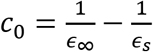 which results in the reduction of the reorganization energy by a factor ≈ 2 when the dielectric constant due to induced dipole *ϵ*_∞_ is accounted for (*ϵ*_s_ is the static dielectric constant). Microscopic simulations and analytical theories^11,12,13^ indicate that the reduction is smaller than suggested by continuum theories, but the problem has not been studied in depth for protein electron transfer.^14^ With *ϵ*_∞_ of water, the reduction compared to nonpolarizable solvents is 0.8 and this factor was used here to rescale the reorganization energies from MD simulations. The corrected λ and *λ*^*St*^ are the MD values (Table S1) multiplied by a factor of 0.8. As the result of this correction, the reaction reorganization energy in Eq 5 is also multiplied by 0.8.

### Rate Calculations

Applying both relaxation times to Eq S1, we find that, due to a large relaxation time of distance fluctuations *τ*_R_ = 281 ps, the term including *τ*_X_/*τ*_R_ in the denominator of Eq. 1 can be dropped. A simpler result for the crossover parameter g follows

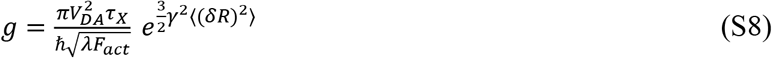

We also find that the parameter g is large, g ≈ 55 at *R* ≈ 0.6nm, when *γ* = 14.4 nm^−1^ and 〈(*δR*)^2^〉 = 0.041^2^ nm^−1^ are adopted. The parameter *γ* is taken from the Hopfield equation (Eq. S2) used to calculate the electronic coupling and the variance of the donor-acceptor distance is from MD simulations of the Y55-Y76 pair adopted for all other donor-acceptor pairs. This value of *g* implies that, at relatively short distances, electron hops between the residues are in the dynamics controlled regime when the rate constant is not affected by the electronic coupling. This happens because the total reaction rate includes the parameter g in the denominator of the following relation

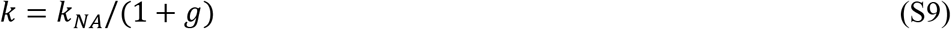

Here, *k*_NA_ is the non-adiabatic rate constant given by the equation

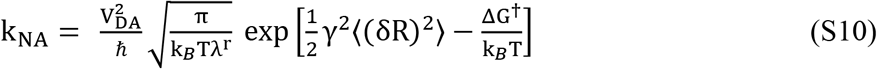

The dynamics-controlled electron-transfer rate constant becomes

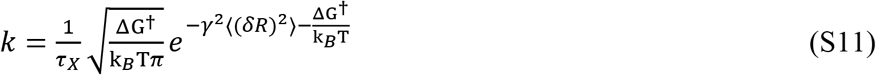

The rate calculations between pairs of residues (Table S1) produce the parameter g spanning a broad range of values such that both the dynamics-controlled and nonadiabtic limits apply to specific intra-protein electron hops. The appearance of the crossover parameter g in the form given by Eqs. S1 and S8 leads to slight deviations from the detailed balance condition requiring the ratio of the forward and backward rate constant to be equal to the Boltzmann factor exp[*−βΔG*]. Given that the Derrida model used here for the calculation of the carriers diffusion constant requires detailed balance, we used a somewhat simplified form of the parameter g replacing ΔG^†^ in Eqs. 1 and 9 with *λ*^*r*^/4. The correction is mostly insignificant for the rate values, but strictly preserves the detailed balance.

### Diffusion constant from the Derrida model

Following Derrida^17^

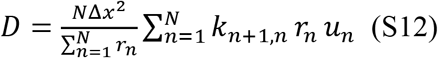

where the step size Δ*x* is calculated as the average distance between the residues and

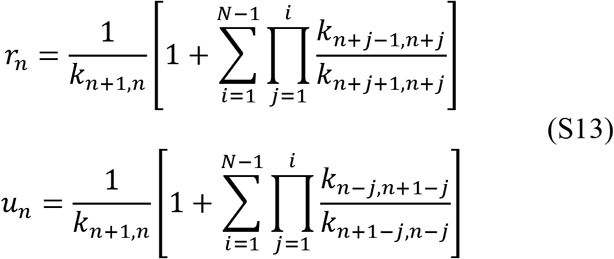

Applied to the path listed in Table S3, these equations yield D = 3.06 nm^2^/ns. Similar numbers are found for other single paths. Applied to the 433 such paths connecting Y36 to Y89, we obtain a sum of the 433 *D* values of 578 nm^2^/ns.

### Charge Injection Calculation

The rate constants for the two contact sites, 1, 2 are

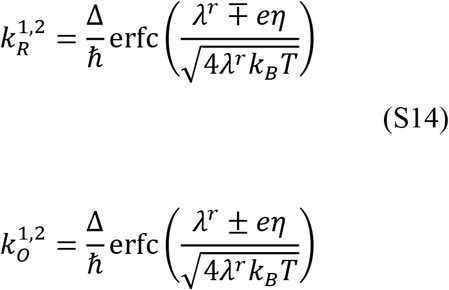

where R and O refer to reduction or oxidation and we assume the same electronic coupling to the contacts, Δ, and the potential drop at the contacts, *η*, is given in the caption to Figure S2. “−“ and “+” refer to 1 and 2, respectively.

The kinetic equation for the fraction of holes *n* at the single injection site is

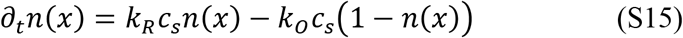

Where 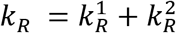 and 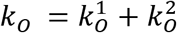 and *c_s_=1/S* is the surface concentration determined by the area of the contact, *S*. At the stationary condition determined by *∂*_t_*n*(*x*) = 0, one gets 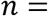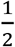 and the following expression for the total current *I* through the contact

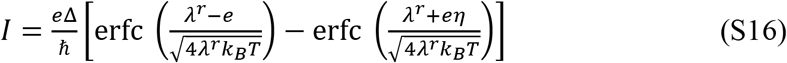

### Kinetic Monte Carlo Model and Brownian diffusion

We perform single charge kinetic Monte Carlo based on a previous routine implemented in python^21^ to simulate charge hopping in the graph based representation of the protein. Each node in the graph is connected by two edges, one for forward and one for backward rates. Every iteration of the Monte Carlo involves determining the lifetime of the charge on the node *i* which it currently resides on and the node which the charge hops to next. The charge can only hop to the site which are connected to its current node by an edge. First, the lifetime of the charge *τ* is determined from an exponential random variable T as τ = T/Γ_i_, where Γ_i_ is the sum of the weights of the edges which originate at node *i*.

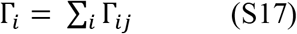

Γ_ij_ is the weight of the edge connecting nodes *i* to *j* which is essentially the rate of charge transfer between the residues represented by *i* and *j*. Next, to determine which residue, the charge hops to, we calculate the probability of hopping to that residue/node *j*.

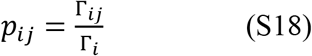

A random number *u* is drawn from a uniform distribution and the next site k is determined from the inequality.

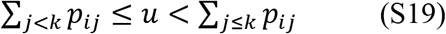

The Monte Carlo (MC) simulation ends when the charge arrives at the residue in the protein which is in contact with the electrodes. We store the residence time τ for every hop and the distance Δ*x*_*i*_ between the center of mass of the residue sidechains travelled during every hop. We then calculate the average Brownian diffusion for that path.

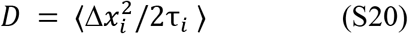

The diffusion constant is calculated for 100000 MC with the starting residue as W35. Another 100000 MC runs are performed for Y36 as the starting residue, summarized Figure 6 of the main text. The resultant diffusion constant values are divided into 300 bins to obtain a distribution, which is fit to a gamma function probability distribution function given by the expression

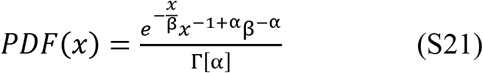

Here α and β are parameters which determine the skewness and peak location of the distribution respectively. We assume β to be the effective diffusion constant for the transport. This gives us 22.13 nm^2^/ns for W35 and 22.8 nm^2^/ns for Y36.

---

**Table S1:** Acceptor (first column) and donor (second column) residue numbers and the reorganization energies calculated from directly from the MD simulations without accounting for the polarization. Also listed (sixth column) the edge-to-edge distance between the residue pairs used in the Hopfield equation. The last column lists the reaction reorganization energies.

| Acceptor | Donor | $\lambda^{St}(eV)$ | $\lambda_{var}(eV)$ | $\Delta G(eV)$ | Edge Distance ($\text{\AA}$) | $\lambda^r$ |
| --- | --- | --- | --- | --- | --- | --- |
| 35 | 36 | 0.55 | 1.422 | 0.18 | 4.51 | 0.213 |
| 35 | 42 | 0.561 | 1.482 | 0.18 | 9.097 | 0.212 |
| 35 | 43 | 0.632 | 1.834 | 0.18 | 11.542 | 0.217 |
| 35 | 54 | 0.543 | 0.881 | 0.18 | 3.437 | 0.335 |
| 35 | 55 | 0.756 | 1.3 | 0.18 | 5.45 | 0.44 |
| 36 | 42 | 0.407 | 1.634 | 0.0 | 9.529 | 0.101 |
| 36 | 43 | 0.514 | 1.3 | 0.0 | 8.49 | 0.203 |
| 36 | 54 | 0.484 | 1.561 | 0.0 | 6.674 | 0.15 |
| 36 | 55 | 0.477 | 0.584 | 0.0 | 2.421 | 0.39 |
| 42 | 43 | 0.456 | 1.249 | 0.0 | 4.85 | 0.166 |
| 42 | 48 | 0.383 | 1.756 | 0.0 | 6.683 | 0.083 |
| 42 | 54 | 0.301 | 0.582 | 0.0 | 7.124 | 0.156 |
| 42 | 55 | 0.492 | 1.253 | 0.0 | 7.069 | 0.193 |
| 43 | 48 | 0.348 | 1.312 | 0.0 | 3.419 | 0.092 |
| 43 | 54 | 0.55 | 1.351 | 0.0 | 8.177 | 0.224 |
| 43 | 55 | 0.411 | 0.617 | 0.0 | 4.136 | 0.274 |
| 48 | 54 | 0.51 | 1.871 | 0.0 | 10.181 | 0.139 |
| 48 | 55 | 0.518 | 1.115 | 0.0 | 5.98 | 0.24 |
| 54 | 55 | 0.542 | 1.232 | 0.0 | 4.737 | 0.239 |
| 36 | 69 | 0.631 | 1.42 | -0.18 | 8.594 | 0.28 |
| 36 | 70 | 0.382 | 1.211 | 0.0 | 7.384 | 0.12 |
| 36 | 88 | 0.542 | 1.589 | 0.0 | 9.799 | 0.185 |
| 43 | 70 | 0.465 | 1.512 | 0.0 | 7.35 | 0.143 |
| 43 | 76 | 0.528 | 1.235 | 0.0 | 7.441 | 0.225 |
| 43 | 77 | 0.524 | 1.15 | 0.0 | 6.308 | 0.239 |
| 43 | 88 | 0.529 | 1.364 | 0.0 | 10.328 | 0.205 |
| 43 | 89 | 0.645 | 1.521 | 0.0 | 6.903 | 0.274 |
| 48 | 70 | 0.328 | 1.529 | 0.0 | 12.525 | 0.07 |
| 48 | 76 | 0.465 | 1.677 | 0.0 | 7.543 | 0.129 |
| 48 | 77 | 0.462 | 1.535 | 0.0 | 8.233 | 0.139 |
| 55 | 69 | 0.809 | 1.3 | -0.18 | 6.901 | 0.503 |
| 55 | 70 | 0.544 | 0.983 | 0.0 | 9.022 | 0.301 |
| 55 | 76 | 0.654 | 1.087 | 0.0 | 6.001 | 0.393 |
| 55 | 77 | 0.45 | 0.964 | 0.0 | 8.64 | 0.21 |
| 55 | 88 | 0.697 | 1.126 | 0.0 | 7.775 | 0.432 |
| 55 | 89 | 0.768 | 0.949 | 0.0 | 8.734 | 0.621 |

**Table S2:**
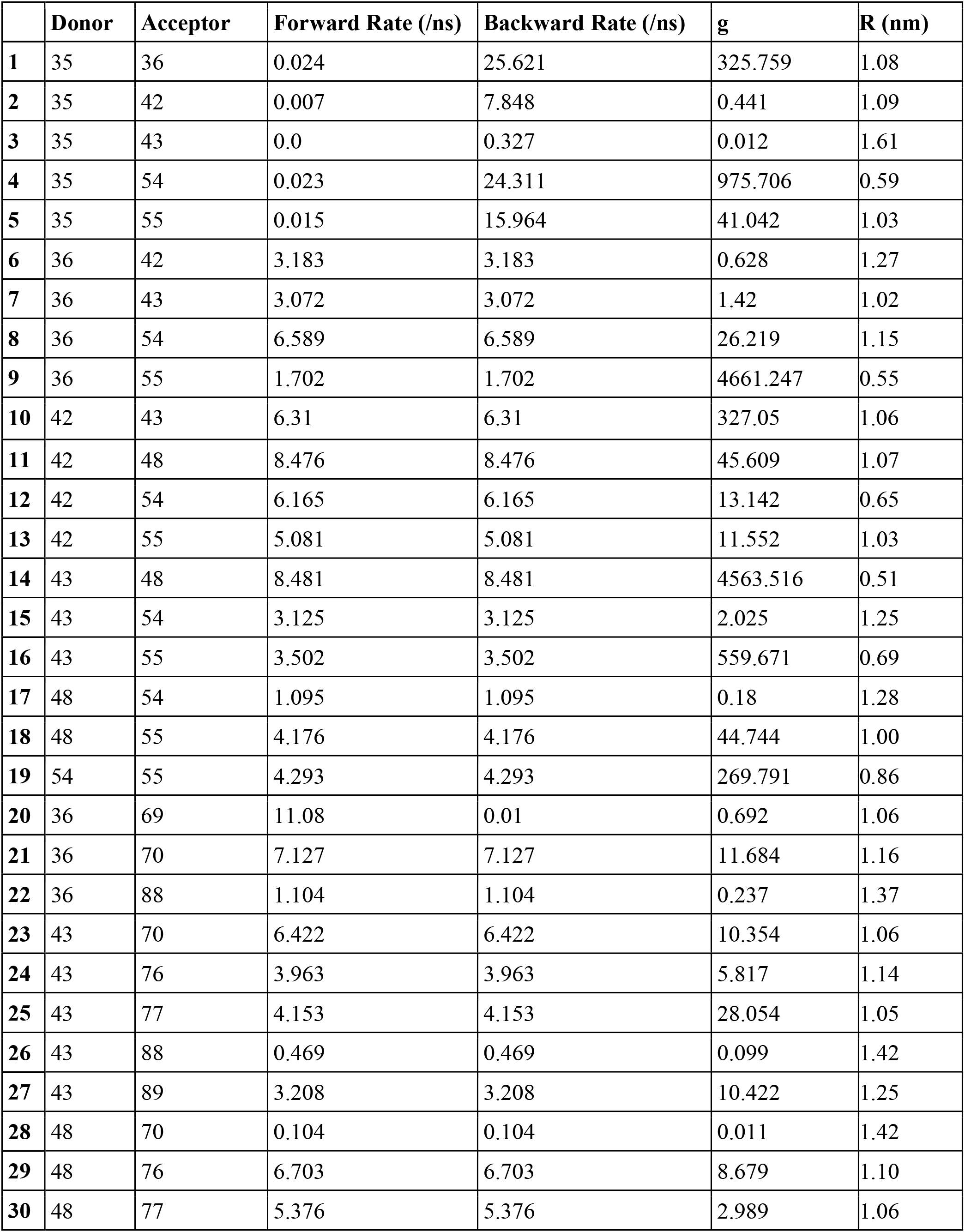
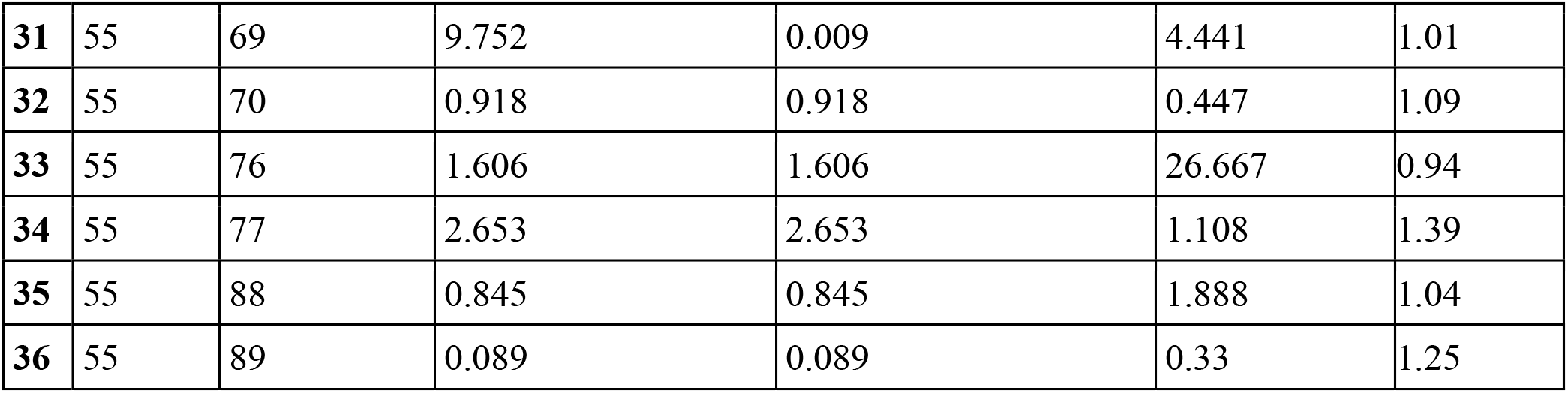
Acceptor (second column) and donor (third column) residue numbers and forward (4^th^ column) and backward rate (5^th^ column). The 6^th^ column lists the crossover parameter g and the last column lists the average COM-COM distance between the residues (averaged over 3000 MD frames). The rates have been calculated with the adjusted reorganization energy to account for the non-polarizable MD forcefield which was used.

|  | **Donor** | **Acceptor** | **Forward Rate (/ns)** | **Backward Rate (/ns)** | **g** | **R (nm)** |
| --- | --- | --- | --- | --- | --- | --- |
| **1** | 35 | 36 | 0.024 | 25.621 | 325.759 | 1.08 |
| **2** | 35 | 42 | 0.007 | 7.848 | 0.441 | 1.09 |
| **3** | 35 | 43 | 0.0 | 0.327 | 0.012 | 1.61 |
| **4** | 35 | 54 | 0.023 | 24.311 | 975.706 | 0.59 |
| **5** | 35 | 55 | 0.015 | 15.964 | 41.042 | 1.03 |
| **6** | 36 | 42 | 3.183 | 3.183 | 0.628 | 1.27 |
| **7** | 36 | 43 | 3.072 | 3.072 | 1.42 | 1.02 |
| **8** | 36 | 54 | 6.589 | 6.589 | 26.219 | 1.15 |
| **9** | 36 | 55 | 1.702 | 1.702 | 4661.247 | 0.55 |
| **10** | 42 | 43 | 6.31 | 6.31 | 327.05 | 1.06 |
| **11** | 42 | 48 | 8.476 | 8.476 | 45.609 | 1.07 |
| **12** | 42 | 54 | 6.165 | 6.165 | 13.142 | 0.65 |
| **13** | 42 | 55 | 5.081 | 5.081 | 11.552 | 1.03 |
| **14** | 43 | 48 | 8.481 | 8.481 | 4563.516 | 0.51 |
| **15** | 43 | 54 | 3.125 | 3.125 | 2.025 | 1.25 |
| **16** | 43 | 55 | 3.502 | 3.502 | 559.671 | 0.69 |
| **17** | 48 | 54 | 1.095 | 1.095 | 0.18 | 1.28 |
| **18** | 48 | 55 | 4.176 | 4.176 | 44.744 | 1.00 |
| **19** | 54 | 55 | 4.293 | 4.293 | 269.791 | 0.86 |
| **20** | 36 | 69 | 11.08 | 0.01 | 0.692 | 1.06 |
| **21** | 36 | 70 | 7.127 | 7.127 | 11.684 | 1.16 |
| **22** | 36 | 88 | 1.104 | 1.104 | 0.237 | 1.37 |
| **23** | 43 | 70 | 6.422 | 6.422 | 10.354 | 1.06 |
| **24** | 43 | 76 | 3.963 | 3.963 | 5.817 | 1.14 |
| **25** | 43 | 77 | 4.153 | 4.153 | 28.054 | 1.05 |
| **26** | 43 | 88 | 0.469 | 0.469 | 0.099 | 1.42 |
| **27** | 43 | 89 | 3.208 | 3.208 | 10.422 | 1.25 |
| **28** | 48 | 70 | 0.104 | 0.104 | 0.011 | 1.42 |
| **29** | 48 | 76 | 6.703 | 6.703 | 8.679 | 1.10 |
| **30** | 48 | 77 | 5.376 | 5.376 | 2.989 | 1.06 |
| **31** | 55 | 69 | 9.752 | 0.009 | 4.441 | 1.01 |
| **32** | 55 | 70 | 0.918 | 0.918 | 0.447 | 1.09 |
| **33** | 55 | 76 | 1.606 | 1.606 | 26.667 | 0.94 |
| **34** | 55 | 77 | 2.653 | 2.653 | 1.108 | 1.39 |
| **35** | 55 | 88 | 0.845 | 0.845 | 1.888 | 1.04 |
| **36** | 55 | 89 | 0.089 | 0.089 | 0.33 | 1.25 |

**Table S3:** Forward and backward rates and the crossover parameter g for transitions in a path from the start of first unit of CTPR8 protein to the last active residue in the second CTPR8 protein. The path was calculated using Dijkstra’s shortest path algorithm^13^. Derrida’s model^16^ is applied to the path to calculate the diffusion constant of 3.06 nm^2^/ns.

| n | Residue Pairs | Forward Rate (ns<sup>-1</sup>) | Backward Rate (ns<sup>-1</sup>) | $g$ |
| --- | --- | --- | --- | --- |
| 1 | 35-36 | 0.024 | 25.621 | 325.759 |
| 2 | 36-43 | 3.072 | 3.072 | 1.42 |
| 3 | 43-89 | 3.208 | 3.208 | 10.422 |
| 4 | 89-35 | 9.752 | 0.009 | 4.441 |

**Table S4:** Partial charges for Y-H^+^ in units of the elementary charge^19^. Atom names are consistent with the CHARMM36 convention. The total sum of the partial charges is +1. The partial charges are applied to the residue and the protein is simulated using NAMD. The trajectory is used to calculate the electrostatic potential map of either the initial or final state during the charge transfer process. The initial state has the electron acceptor residue positively charged while the final state has the donor residue positively charged.

| Atom | Charge ($e$) |
| --- | --- |
| CB | -0.18000000715255737 |
| HB1 | 0.09000000357627869 |
| HB2 | 0.09000000357627869 |
| CG | 0.42800000309944153 |
| CD1 | -0.23000000417232513 |
| HD1 | 0.22300000488758087 |
| CE1 | -0.23000000417232513 |
| HE1 | 0.22300000488758087 |
| CZ | 0.6209999918937683 |
| OH | -0.4300000071525574 |
| HH | 0.4090000092983246 |
| CD2 | -0.23000000417232513 |
| HD2 | 0.22300000488758087 |
| CE2 | -0.23000000417232513 |
| HE2 | 0.11500000208616257 |

**Table S5:** Partial charges for W-H^.+^ in units of the elementary charge^20^. Atom names are consistent with the CHARMM36 convention. The total sum of the partial charges is +1. The partial charges are applied to the residue and the protein is simulated using NAMD. The trajectory is used to calculate the electrostatic potential map of either the initial or final state during the charge transfer process. The initial state has the electron acceptor residue positively charged while the final state has the donor residue positively charged.

| Atom | Charge ($e$) |
| --- | --- |
| CB | 0.07000000029802322 |
| HB1 | 0.009999999776482582 |
| HB2 | 0.009999999776482582 |
| CG | 0.11999999731779099 |
| CD1 | 0.12999999523162842 |
| HD1 | 0.004999999888241291 |
| NE1 | -0.14000000059604645 |
| HE1 | 0.4699999988079071 |
| CE2 | -0.009999999776482582 |
| CD2 | -0.07000000029802322 |
| CE3 | 0.029999999329447746 |
| HE3 | 0.004999999888241291 |
| CZ3 | 0.15000000596046448 |
| HZ3 | 0.029999999329447746 |
| CZ2 | 0.12999999523162842 |
| HZ2 | 0.004999999888241291 |
| CH2 | 0.05000000074505806 |
| HH2 | 0.004999999888241291 |

